# Cardiometabolic Recovery and Lactate Removal may be Related to Muscular Adaptations

**DOI:** 10.1101/402511

**Authors:** Edgar Ismael Alarcón Meza, Glauber Lameira de Oliveira, Talita Adão Perini de Oliveira, Marcelo Hubner Moreira, Angeliete Garcez Militão, José Fernandes Filho, Vernon Furtado Da Silva, Estélio Henrique Martin Dantas, João Rafael Valentim-Silva

## Abstract

The aim of this study was to measure the differences in cardiorespiratory recovery (CR) and blood lactate removal among young athletes with differences in non-lactic (NP) and lactic anaerobic power (LP) and fatigue index (FI) but with the same degree of cardiorespiratory fitness. Sixteen swimmers from the Brazilian synchronized swimming team (2014) were divided into two groups GAP (Group High Power) (n = 9), with NP, LP, and FI (p<0.05) compared to GBP (Group Low Power) (n=7). Both groups performed a four-minute routine at competitive intensity. Anaerobic power, maximal heart rate (HR) and blood lactate (BL) were determined before and at 1, 3 and 5 minutes after the routine. Student’s t-test was used to analyze the intergroup differences of NP, LP, FI, maximum and lactic HR, and two-way ANOVA followed by Bonferroni was used to analyze HR and BL at 1, 3 and 5 minutes after activity with a significance of 5%. The FI of the GBP group was lower than that of the GAP group (P <0.05). The NP of the GBP group was higher than that of the GAP group (P <0.05). The maximum HR of the GBP group was equal to that of the GAP group (P> 0.05). The GBP group had better HR recovery than did the GAP group (P <0.05). BL had its lowest levels after 1 and 5 minutes of recovery in the GBP group when compared to the GAP group (P <0.05). The GBP group’s FI was significantly lower than that of the GAP group, while NP was higher, and CR was better in the GBP group, indicating a relationship between a lower FI and higher NP and LP with CR and suggesting that muscular adaptations have an important influence on CR and BL removal.

## Introduction

The literature is consistent in affirming that various psychological, biological and performance variables are indicators of fatigue. Investigations have revealed that creatine kinase, lactate dehydrogenase, blood lactate, oxygen consumption and heart rate are associated with exercise intensity and fatigue. Furthermore, the time required to return to pre-exercise levels is considered an indicator of the recovery capacity that is linked to an athlete’s physical fitness [1–4].

Heart rate variability (HRV) analysis is an established method used to quantify the extent of autonomic recovery from exercise [5,6]. After completing a given exercise, a rapid decrease in parasympathetic cardiac activity to resting levels suggests a relative and physiological systemic recovery imposed by the workload [7,8] and the amount of time required for parasympathetic reactivation after exercise may be significantly influenced by several factors, including exercise intensity[9] and cardiorespiratory fitness [10,11]. Although many of the factors inherent to HRV have been frequently studied, studies specifically concerning the non-lactic and lactic anaerobic capacities and fatigue index have not yet been conducted.

Another important marker of metabolic activation provided by intense exercise is lactate. Many authors have shown that when the rate of ATP production by oxidative sources becomes insufficient, high rates of glycolytic or glycogenolytic ATP production are required, culminating in the production of pyruvate, a metabolite that can be reduced to lactate or oxidized to CO_2_ or H_2_O [12–15].

Thus, by increasing the intensity of exercise and muscle load, several tissues begin to produce more lactate and export it to the circulation. Simultaneously, the less active skeletal muscles, the heart, the liver, the renal cortex and the brain remove lactate from circulation, suggesting that this metabolite acts as an intermediary for transporting carbohydrates from cells and tissues with relatively low oxidative capacity to cells and tissues with high oxidative capacity [16,17]. Therefore, it is well established that blood lactate concentration is the result of production and removal of this metabolite.

Lactate has been a focus of research on skeletal muscle metabolism and a central theme of the controversy over the relationships between lactate production, acidosis and fatigue, because it is probably a residual product; lactate is increasingly regarded as a source of fuel, continuously produced and metabolized in the organism in order to maintain energy homeostasis at adequate levels in an attempt to maintain cellular activity [12,16,18].

A fact that somewhat contradicts what has been described in the literature up to the present moment is that high-intensity exercise can produce greater expression and translocation of MCT-4 (monocarboxylate transporter) [19,20]. This adaptation may have implications for the export and import of lactate, because the plasma membrane is responsible for this transport; furthermore, high-intensity exercise, in addition to promoting adaptation related to lactic and non-lactic anaerobic resistance, is the stimulus for MCT-4 translocation and expression in muscles [21,22].

These statements support the notion that muscular fitness levels may be related to reduced blood lactate levels, while cardiorespiratory fitness is seen as being analogous to the velocity of decrease in heart rate, which is the hypothesis postulated in the present investigation. Thus, the objective of the present study is based on the possibility that athletes with various levels of non-lactic and lactic power and fatigue indexes present various behaviors in terms of lactate removal and heart rate recovery curves.

## Materials and Methods

### Volunteer Group

All 16 swimmers (17 ± 1.4 years old) from the Brazilian synchronized swimming team (2014) participated in the study at the beginning of the training season (January). The athletes were divided into two groups – GAP (n= 9) and GBP (n= 7) – based on their non-lactic and lactic anaerobic capacity. The athletes were instructed not to train the day before the test, to have their last meal up to three hours ahead of time, to avoid caffeine intake, to avoid medication, and to sleep at least 6 hours the night before. A brief history was taken to identify any possible exclusion factors. The volunteer characteristics are summarized in the Table 1.

**Table 1:** physical data and training experience of the volunteers

| Variables | GAP (n=9) | GBP (n=7) |
| --- | --- | --- |
| Age (years) | 18.8±1.4 | 15.8± 1.6 |
| Height (cm) | 166.31±4.3 | 162.0±3.1 |
| Body Mass | 55.16±4.2 | 55.28±5.2 |
| Body Mass Index | 20.01±3.3 | 21.06±3.7 |
| Training Time | 9.5±1.6 | 7.9±1.7 |

Values are expressed as the mean ± SD.

### Procedures

#### Exercise Protocol

The exercises were composed of a competitive routine with a duration of 4 minutes at maximum competitive intensity. Prior to the tests, heart rates and lactate levels were measured. Then, the competitive routine was performed with heart rate monitoring in order to determine maximum heart rate. Finally, after 1, 3 and 5 minutes of recovery, heart rate and blood samples were collected for later determination of blood lactate levels as summarized in figure 1.

**Figure 1:**
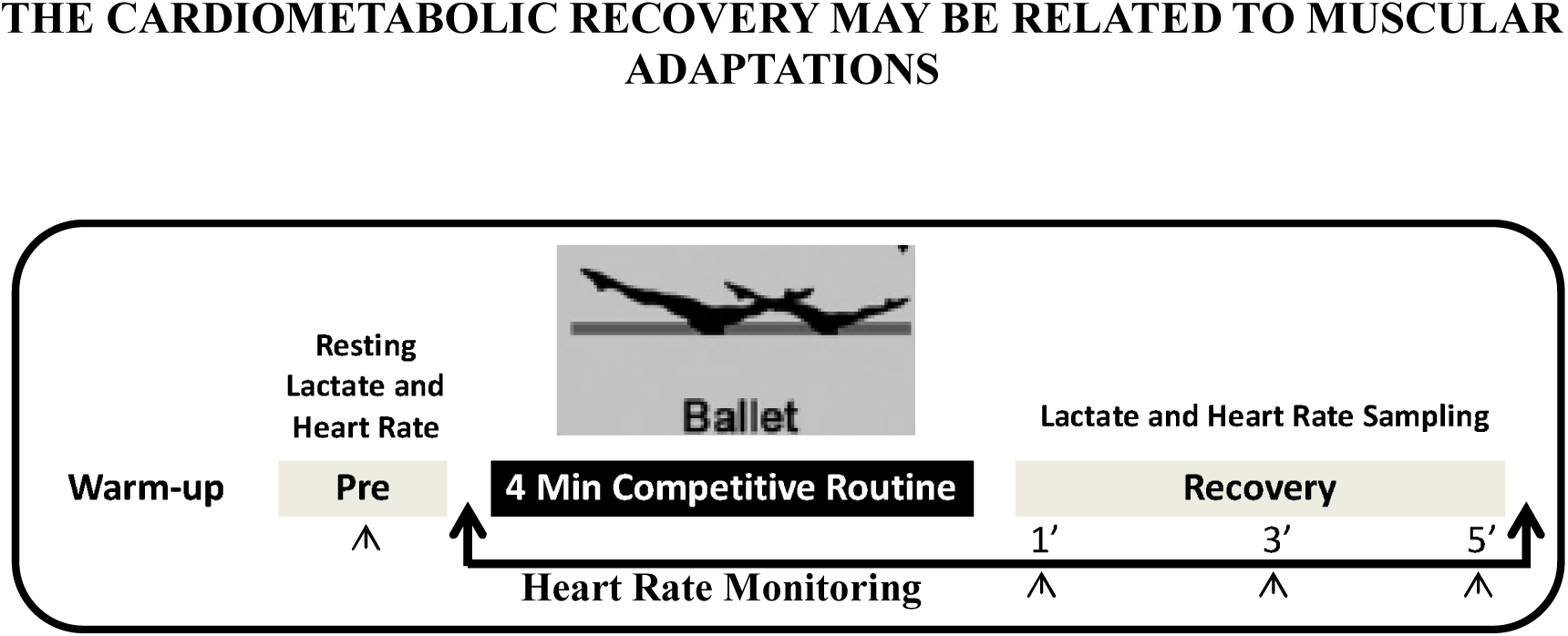
exercise protocol and data collection.

#### Functional Capacity

The stress test was performed on a treadmill (ECAFIX), using Bruce’s protocol. The Borg scale was adopted for a subjective measure of effort. The test was interrupted in the presence of symptoms that impeded its continuity (fatigue) and/or represented a risk to the evaluated individual. The exercise period was preceded by a period of adaptation and preparation of the athlete to the equipment, followed by five minutes of monitored active recovery (five km/h).

Gas exchange was measured with a pneumotachograph (Pt, MEDGRAFIC, Wilmington, DE, USA) coupled to a differential pressure transducer, near the capillary of the gas analyzer in order to collect a sample of inhaled and exhaled gases. The athlete, whose nose was sealed by a clip, was connected to the system for acquisition of ventilatory data through a mouthpiece. Flow signals and gaseous concentrations were sampled (Pentium III computer) at a rate of 1000 Hz. Cycle-to-cycle gas concentrations were sampled by the cardiorespiratory diagnostic system MEDGRAFIC, model VO2000. Respiratory flow and electrocardiographic signals (ECG - ECAFIX) were processed on a personal computer in real time.

The mean values of the maximum load reached during exercise of the last three respiratory cycles were computed in order to determine the results of the variables representative of the end of the exercise (peak). The following ventilatory variables were analyzed: ventilation minute (V_E_, l.min^-1^ – BTPS), oxygen consumption (VO_2peak_, l.min^-1^ – STPD), carbon dioxide production (VCO_2peak_, l.min^-1^ – STPD), gas exchange ratio (R=VCO_2peak_/VO_2peak_) and ventilatory anaerobic threshold (LA, 1.min^-1^ – STPD).

Identification of LA was done through a non-invasive method from the analysis of the ventilatory equivalent of oxygen (V_E_/VO_2_) and carbon dioxide (V_E_/VCO_2_), as previously described [23,24].

The gas analyzer was calibrated daily before beginning the tests with a calibration bullet (AGA – primary standard) with the following concentrations: 12.1% O_2_, 5.0% CO_2_ and 83.0% N_2_.

#### Anaerobic Power

Non-lactic and lactic anaerobic power and the fatigue rate were determined by the Wingate test [25] performed on a mechanical cycle ergometer (MONARK). This test preceded the ergospirometric test with an interval of at least 60 minutes between them. To compute the total revolution of the pedal and to calculate the power every five seconds, an optical sensor was coupled to the cycle ergometer managed by an acquisition program elaborated in Labview 6.0 (National Instruments, USA).

The load was determined by 0.075 Kp.kg^-1^ of body mass as predicted in the Wingate test protocol. The test was preceded by a two-minute warm-up period and followed by an active three-minute recovery period with a 50-watt load. From the individual results, the following parameters were calculated: non-lactic anaerobic power by means of the absolute (or maximum) power peak (watts) and relative to body mass (watts.kg^1^) and the lactic power or average (watts). The percentage of power decline (%) or rate of fatigue was obtained from the difference between the highest power and the lowest power achieved by the athlete during the 30 s test. The bicycle load was checked before starting the test.

Parameter calculations, descriptive statistics of the data and comparisons were performed using one-way ANOVA and the Bonferroni post-tests in the program “PrismStat 5.0” with a significance level of p <0.05.

#### Ethical aspects

The evaluations were carried out in the Ergo-Spirometry Department of the Laboratory of Exercise Physiology of the School of Physical Education and Sports at the Federal University of Rio de Janeiro (SE-LABOFISE-EEFD-UFRJ).

All participants were aware and signed a written consent form, including the procedures adopted and the authorization of the volunteers to investigate the results found in the scientific study. The anonymity and privacy of the participants were preserved in the study. The Council of Ethics and Human Research of the Federal Institute of Education of Rondônia approved this investigation under CAAE number 44907715.2.00005653. All subjects gave their consent to participate.

## Results

Heart rate recovered faster in the GBP group than in the GAP group (P<0.05) (Fig. 2A); a similar phenomenon was observed regarding the behavior of lactate (P <0.05) (Fig. 2B). In relation to physical fitness and respiratory capacities, there were no differences between the groups (P> 0.05) (Fig. 2C and 2D); however, the fatigue index, lactic and non-lactic anaerobic power showed substantial differences between the groups (P <0.0001) (Fig. 2E).

**Figure 2:**
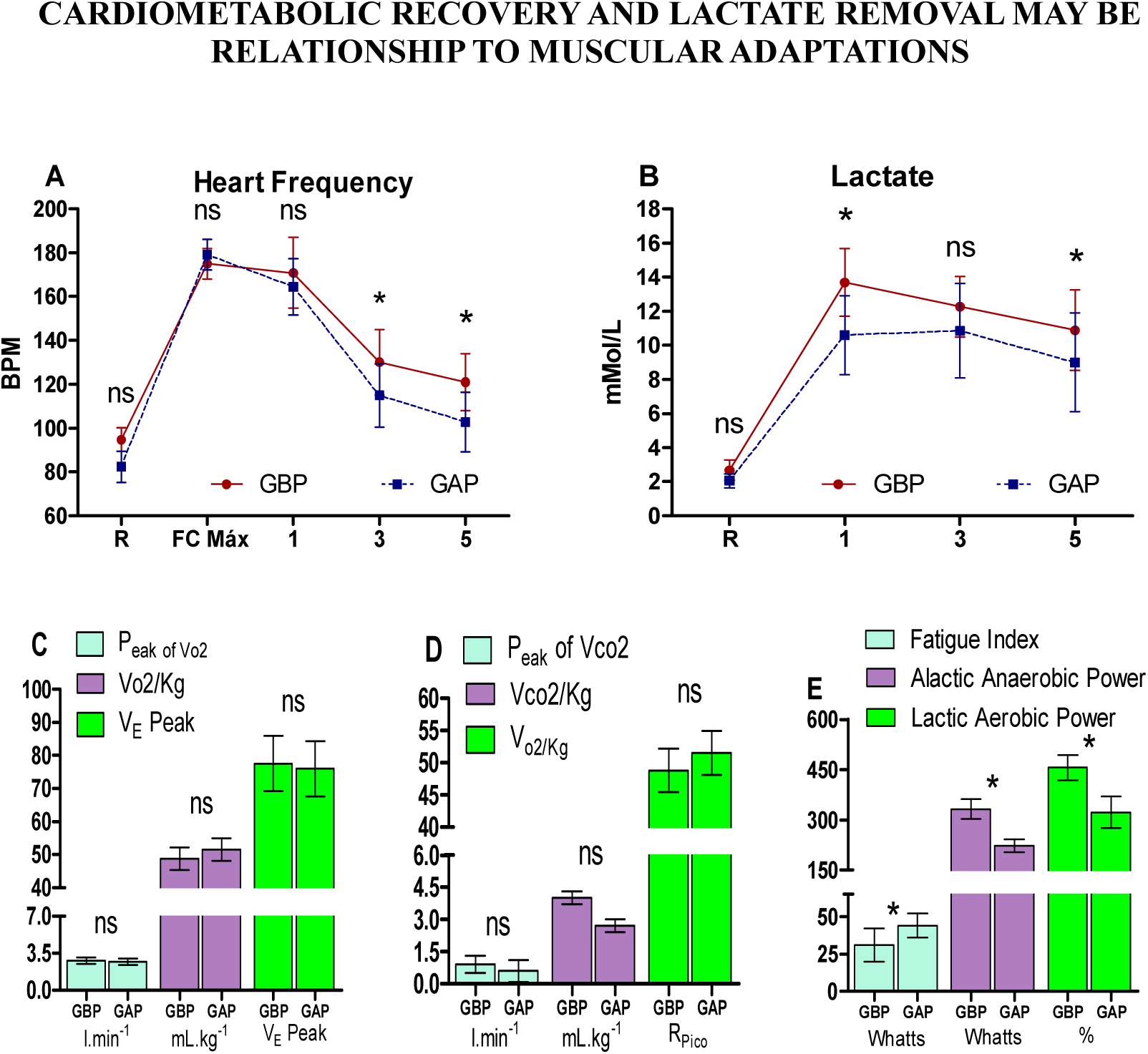
heart rate and lactate recovery curve. The two groups, GBP (n=9) and GAP (n=7), were subjected to a 4-minute routine on different days. **(A)** Comparison of heart rates between the GBP and GAP groups before, during and after the 4-minute routine. **(B)** Comparison of lactate between the GBP and GAP groups before, during and after the 4-minute routine. **(C)** Comparison of the VO_2_, VO_2_/kg peak, and the V_E_ peak between the GBP and GAP groups. **(D)** Comparison of the R Peak, VO_2_ and VO_2_/kg peak between the GBP and GAP groups. (E). Comparison of the Fatigue Index, Non-lactic Anaerobic Power and Lactic Anaerobic Power between the GBP and GAP groups **(A, B, C** and **D** ns = p> 0.05, *p <0.05). **(G** * = p <0.0001 GBP vs GAP).

## Discussion

The present study is the first in the literature to compare the cardiometabolic responses of elite athletes of synchronized swimming in terms of non-lactic anaerobic, lactic anaerobic power and fatigue index. To determine the physical characteristics of both groups, some measures of body composition, functional capacity and anaerobic power were conducted. Recently, it was shown that lactate removal was partly dependent on the time of recovery between stimulus sessions, and moderate correlations were observed between post-exercise blood lactate concentration, peak heart rate and perceived exertion [26]. Nevertheless, the possible physiological mechanisms underlying these observations were not described. Here, we propose that the muscle adaptations regarding the fatigue index can be related to lactate removal and heart hate recovery.

It is well established in the literature that lower fatigue index and higher non-lactic and lactic anaerobic power are associated with higher level of exercise-related muscle adaptations. It has been proposed that aerobic training may improve the ability of muscle to recover after anaerobic exercise, suggesting that an athlete with greater aerobic fitness will use fewer non-oxidative sources and thus will recover more quickly from exercise. Theoretically, an increase in aerobic fitness improves the recovery from anaerobic exercise However, this phenomenon was not completely reproduced in the present study because the aerobic capacity of both groups was equal while the cardiometabolic recovery was different. This is reasonable because lactate removal could be linked to uptake by muscle and other tissues that use this substrate as an energy source [12,27].

A synchronized swimming routine requires a high aerobic effort with fundamental anaerobic contributions to maintain adequate exercise levels during competition [28,29]. The data shown here demonstrate that the peaks of non-lactic and lactic anaerobic power have the same behavior, but the fatigue index was the only factor directly related to physical fitness that showed differences between the groups.

In the present study, the GAP group had heart rate and lactate curve response behaviors different from those observed in the GBP group, although the maximum heart rate was equal in both groups. The GBP group showed a lower lactate peak than did the GAP group. Another interesting fact is that the non-lactic anaerobic power, lactic anaerobic power and the fatigue index differed between the two groups, even though VO_2_ and VCO_2_ were equal. These data contradict a part of the literature that states that aerobic capacity improves recovery after intense exercise, a finding that was not seen in the present investigation.

Other investigations have pointed to several possible explanations for the effect of aerobic conditioning on recovery. Holloszy [30] pointed to the increase in the number and size of mitochondria in metabolic recovery. However, the increased myoglobin content and blood volume together would be implicated in the transport of O_2_ to the muscle during high-intensity exercise, and consequently, there would be a decrease in recovery time, since this would also help to remove the accumulated lactate more quickly [31,32].

These data explain the higher velocity in cardiometabolic recovery, However, the data described here demonstrate a clear difference in lactate removal between the groups though both exhibit equal V_E_ peak, V_O2 Peak_, V_O2 Peak/body mass_ and peak heart rate, suggesting that the aerobic capacity of these athletes is unrelated to metabolic recovery after maximal intensity exercise.

This difference may be related to the use of lactate by other muscles and tissues, since anaerobic metabolism after a few minutes of intense exercise may activate lactate uptake and use it as an important energy source during intense exercise by modifying the kinetics of the use of glucose in the presence of lactate[15]. However, unfortunately, almost all studies investigating differences in lactate responses depend on blood lactate measurements that reflect muscle lactate alone, providing indirect evidence of lactate accumulation and removal. Other authors did not find evidence that associated the aerobic capacity with greater speed of lactate removal, corroborating the findings of the present study [32–34]. These data, taken together, suggest that the adaptations that led to differences in lactate responses, but not in heart rate, were linked to the muscular system in greater magnitude than to the cardiorespiratory system, since the ventilatory capacities between the groups (V_E Peak_, V_O2 Peak_, V_O2 Peak/body mass_, V_CO2 Peak_, V_CO2 Peak/body mass_) were the same. Therefore, it appears that the cardiorespiratory adaptations of both groups reached the same levels, suggesting that the observed difference was related to adaptations of the muscles required during the exercise routine, although explaining the mechanism that is linked to what was observed here was not the focus of the present investigation.

A study conducted by Otsuki et al. [35] demonstrated differences in cardiac recovery rates between athletes trained in endurance and strength. This study may have its results extrapolated to the data found here, because the non-lactic and lactic anaerobic power and the fatigue index between the two groups showed differences, a fact that is usually evident among people who are trained or not in exercises and strength. However, studies that demonstrate differences in heart rate between groups that have differences in non-lactic and lactic anaerobic power and the fatigue index are scarce, limiting the discussion of the present study.

Among the limitations found in the present study is the need for a population with greater diversity than the one tested here and in previous studies; for example, diversity in terms of age, BMI and health status is the next logical step in order to refine the interpretation of the data described here. There is also a scarcity of data regarding metabolic and cardiac recovery in elite athletes with the same level of aerobic conditioning for comparison between our findings and those of other authors.

## Conclusions

Lactate removal and HR characteristics showed differences between the groups, with the best performance and the group with the lowest fatigue index, although cardiorespiratory fitness, demonstrated by the maximum HR reached, and the measure of pulmonary capacity and O_2_ uptake were equal between the two groups. These data suggest that muscular adaptations may have a critical role in the behavior of cardiometabolic recovery, in contrast to the findings in the literature that focused on muscle metabolism in exercise.

## Practical Applications

This paper challenges the idea that only the cardiovascular training is related to recovery after an exercise training session. Therefore, an intense neuromuscular training could help in recovery, especially in sports in which there is more than one competition on the same day. This may be applicable to athletes in sports with fundamentally aerobic components. This knowledge could be crucial for improving performance during the several trials during the competition day.

## Acknowledgments

The authors wish to thank the athletes who participated in the study for their effort and dedication for their help and availability.

---

**Table 1:** volunteer group characterization.

| Variables | GAP (n=9) | GBP (n=7) |
| --- | --- | --- |
| Age (years) | 18.8±1.4 | 15.8± 1.6 |
| Height (cm) | 166.31±4.3 | 162.0±3.1 |
| Body Mass | 55,16±4,2 | 55,28±5,2 |
| Body Mass Index | 20,01±3,3 | 21,06±3,7 |
| Training Time | 9.5±1.6 | 7.9±1.7 |
| Values are expressed as mean ± SD. |  |  |

## References

1. Rider BC, Coughlin AM, Hew-Butler TD, Goslin BR. Effect of compression stockings on physiological responses and running performance in division III collegiate cross-country runners during a maximal treadmill test. J Strength Cond Res. 2014;28(6):1732–1738. doi:10.1519/JSC.0000000000000287.

2. Elias GP, Wyckelsma VL, Varley MC, McKenna MJ, Aughey RJ. Effectiveness of water immersion on postmatch recovery in elite professional footballers. Int J Sports Physiol Perform. 2013;8(3):243–253. doi:10.1123/ijspp.8.3.243.

3. Higgins TR, Climstein M, Cameron M. Evaluation of hydrotherapy, using passive tests and power tests, for recovery across a cyclic week of competitive rugby union. J Strength Cond Res. 2013;27(4):954–965. doi:10.1519/JSC.0b013e318260ed9b.

4. Higgins T, Cameron M, Climstein M. Evaluation of passive recovery, cold water immersion, and contrast baths for recovery, as measured by game performances markers, between two simulated games of rugby union. J Strength Cond Res. 2012:1. doi:10.1519/JSC.0b013e31825c32b9.

5. Michael S, Graham KS, Davis GM. Cardiac Autonomic Responses during Exercise and Post-exercise Recovery Using Heart Rate Variability and Systolic Time Intervals—A Review. Front Physiol. 2017;8. doi:10.3389/fphys.2017.00301.

6. Molina GE, Fontana KE, Porto LGG, Junqueira LF. Post-exercise heart-rate recovery correlates to resting heart-rate variability in healthy men. Clin Auton Res. 2016;26(6):415–421. doi:10.1007/s10286-016-0378-2.

7. Cinaz B, Arnrich B, La Marca R, Tröster G. Monitoring of mental workload levels during an everyday life office-work scenario. Pers Ubiquitous Comput. 2013;17(2):229–239. doi:10.1007/s00779-011-0466-1.

8. Buchheit M, Chivot A, Parouty J, et al. Monitoring endurance running performance using cardiac parasympathetic function. Eur J Appl Physiol. 2010;108(6):1153–1167. doi:10.1007/s00421-009-1317-x.

9. Stuckey MI, Tordi N, Mourot L, et al. Autonomic recovery following sprint interval exercise. Scand J Med Sci Sports. 2012;22(6):756–763. doi:10.1111/j.1600-0838.2011.01320.x.

10. Sandercock G, Hurtado V, Cardoso F. Changes in cardiorespiratory fitness in cardiac rehabilitation patients: A meta-analysis. Int J Cardiol. 2013;167(3):894–902. doi:10.1016/j.ijcard.2011.11.068.

11. Da Silva DF, Bianchini JAA, Antonini VDS, et al. Parasympathetic cardiac activity is associated with cardiorespiratory fitness in overweight and obese adolescents. Pediatr Cardiol. 2014;35(4):684–690. doi:10.1007/s00246-013-0838-6.

12. Brooks GA. Cell-cell and intracellular lactate shuttles. J Physiol. 2009;587(23):5591–5600. doi:10.1113/jphysiol.2009.178350.

13. Brooks GA. Lactate shuttles in nature. Biochem Soc Trans. 2002;30(2):258–264. doi:10.1042/.

14. Brooks GA. Current concepts in lactate exchange. Med Sci Sports Exerc. 1991;23(8):895–906. doi:10.1249/00005768-199108000-00003.

15. Brooks GA. Lactate production under fully aerobic conditions: the lactate shuttle during rest and exercise. Fed Proc. 1986;45(13):2924–2929. http://www.ncbi.nlm.nih.gov/pubmed/3536591.

16. van Hall G, Lundby C, Araoz M, Calbet JAL, Sander M, Saltin B. The lactate paradox revisited in lowlanders during acclimatization to 4100 m and in high-altitude natives. J Physiol. 2009;587(5):1117–1129. doi:10.1113/jphysiol.2008.160846.

17. van Hall G, Strømstad M, Rasmussen P, et al. Blood lactate is an important energy source for the human brain. J Cereb Blood Flow Metab. 2009;29(6):1121–1129. doi:10.1038/jcbfm.2009.35.

18. Faude O, Kindermann W, Meyer T. Lactate threshold concepts: How valid are they? Sport Med. 2009;39(6):469–490. doi:10.2165/00007256-200939060-00003.

19. Mogi A, Koga K, Aoki M, et al. Expression and role of GLUT-1, MCT-1, and MCT-4 in malignant pleural mesothelioma. Virchows Arch. 2013;462(1):83–93. doi:10.1007/s00428-012-1344-6.

20. Luo F, Zou Z, Liu X, et al. Enhanced glycolysis, regulated by HIF-1a via MCT-4, promotes inflammation in arsenite-induced carcinogenesis. Carcinogenesis. 2017;38(6):615–626. doi:10.1093/carcin/bgx034.

21. Ideno M, Kobayashi M, Sasaki S, et al. Involvement of monocarboxylate transporter 1 (SLC16A1) in the uptake of l -lactate in human astrocytes. Life Sci. 2017. doi:10.1016/j.lfs.2017.10.022.

22. Takimoto T, Tsue H, Takahashi H, Tamura R, Sasaki H. Synthesis of p-tert-butylcalix[4]thiacrowns exhibiting sulfur number-dependent complexation with mercury(II) ion. Heterocycles. 2014;88(2):911–917. doi:10.3987/COM-13-S(S)61.

23. Beaver WL, Wasserman K, Whipp BJ. A new method for detecting anaerobic threshold by gas exchange. J Appl Physiol. 1986;60(6):2020–2027. http://www.ncbi.nlm.nih.gov/pubmed/3087938.

24. Dickstein K, Barvik S, Aarsland T, Snapinn S, Karlsson J. A comparison of methodologies in detection of the anaerobic threshold. Circulation. 1989;81(1 Suppl):II38–II46. http://eutils.ncbi.nlm.nih.gov/entrez/eutils/elink.fcgi?dbfrom=pubmed&id=2295151&retmode=ref&cmd=prlinks%5Cnpapers2://publication/uuid/C7D07161-D4E6-4953-A5AD-12B432B33AD8.

25. Bar-Or O. The Wingate Anaerobic Test An Update on Methodology, Reliability and Validity. Sport Med An Int J Appl Med Sci Sport Exerc. 1987;4(6):381–394. doi:10.2165/00007256-198704060-00001.

26. Lessard SJ, Rivas DA, Alves-Wagner AB, et al. Resistance to aerobic exercise training causes metabolic dysfunction and reveals novel exercise-regulated signaling networks. Diabetes. 2013;62(8):2717–2727. doi:10.2337/db13-0062.

27. Rotstein A, Dotan R, Bar-Or O, Tenenbaum G. Effect of training on anaerobic threshold, maximal aerobic power and anaerobic performance of preadolescent boys. Int J Sports Med. 1986;7(5):281–286. doi:10.1055/s-2008-1025775.

28. Yamamura C, Zushi S, Takata K, Ishiko T, Matsui N, Kitagawa K. Physiological characteristics of well-trained synchronized swimmers in relation to performance scores. Int J Sports Med. 1999;20(4):246–251. doi:10.1055/s-2007-971125.

29. Dos-Santos JW, de Mello MA. Responses of Blood Lactate Concentration in Aerobic and Anaerobic Training Protocols at Different Swimming Exercise Intensities in Rats. J Exerc Physiol Online. 2011;14(3):34–42. http://proxy.lib.ohio-state.edu/login?url=http://search.ebscohost.com/login.aspx?direct=true&db=s3h&AN=65237273&site=ehost-live.

30. Holloszy JO, Coyle EF. Adaptations of skeletal muscle to endurance exercise and their metabolic consequences. J Appl Physiol. 1984;56(4):831–838.

31. Kanda K, Sugama K, Hayashida H, et al. Eccentric exercise-induced delayed-onset muscle soreness and changes in markers of muscle damage and inflammation. Exerc Immunol Rev. 2013;19:72–85.

32. Tomlin DL, Wenger H a. The relationship between aerobic fitness and recovery from high intensity intermittent exercise. Sport Med. 2001;31(1):1–11. doi:10.2165/00007256-200131010-00001.

33. Freund H, Lonsdorfer J, Oyono-Enguéllé S, Lonsdorfer A, Bogui P. Lactate exchange and removal abilities in sickle cell patients and in untrained and trained healthy humans. J Appl Physiol. 1992;73(6):2580–2587. http://www.ncbi.nlm.nih.gov/pubmed/1490972.

34. Bassett DR, Merrill PW, Nagle FJ, Agre JC, Sampedro R. Rate of decline in blood lactate after cycling exercise in endurance-trained and -untrained subjects. J Appl Physiol. 1991;70(4):1816–1820. http://www.ncbi.nlm.nih.gov/pubmed/2055859.

35. Otsuki T, Maeda S, Iemitsu M, et al. Postexercise heart rate recovery accelerates in strength-trained athletes. Med Sci Sports Exerc. 2007;39(2):365–370. doi:10.1249/01.mss.0000241647.13220.4c.

